# Construction of light-activated neurotrophin receptors using the improved Light-Induced Dimerizer (iLID)

**DOI:** 10.1101/850412

**Authors:** Jen M. Hope, Aofei Liu, Ghawayne J. Calvin, Bianxiao Cui

## Abstract

Receptor tyrosine kinases (RTKs) play crucial roles in human health, and their misregulation is implicated in disorders ranging from neurodegenerative disorders to cancers. The highly conserved mechanism of activation of RTKs makes them especially appealing candidates for control via optogenetic dimerization methods. This work offers a strategy for using the improved Light-Induced Dimer (iLID) system with a constructed tandem-dimer of its binding partner nano (tdnano) to build light-activatable versions of RTKs. In the absence of light, the iLID-RTK is cytosolic, monomeric and inactive. Under blue light, the iLID + tdnano system recruits two copies of iLID-RTK to tdnano, dimerizing and activating the RTK. We demonstrate that iLID opto-iTrkA and opto-iTrkB are capable of reproducing downstream ERK and Akt signaling only in the presence of tdnano. We further show with our opto-iTrkA that the system is compatible with multi-day and population-level activation of TrkA in PC12 cells. By leveraging genetic targeting of tdnano, we achieve RTK activation at a specific subcellular location even with whole-cell illumination, allowing us to confidently probe the impact of context on signaling outcome.

## Introduction

Receptor tyrosine kinases (RTKs) play a critical role in the regulation of cell survival, proliferation, and differentiation. Over 50 RTKs from 20 different subfamilies have been identified in humans, and their misregulation has been implicated in several disorders including cancers and neurodegenerative diseases. The structure of RTKs is highly conserved: they consist of a ligand-binding extracellular domain (ECD), a single transmembrane helix (TM), and an intracellular kinase domain (ICD) responsible for propagating signal to downstream cascades^1^. While there are some variations among subfamilies, the general mechanism of activation of receptor tyrosine kinases is as follows: ligand binding stabilizes the formation of a dimer, or sometimes oligomer, of the RTK; *trans*-autophosphorylation occurs in the activation loop of the ICD, destabilizing the *cis*-autoinhibition of the kinase; and the kinase *trans*-autophosphorylates other tyrosines in the ICD which serve as docking sites for downstream signaling molecules^2^.

Our group and others have successfully leveraged the *Arabidopsis thaliana* photoreceptor CRY2^3^ to make light-inducible constructs of several RTKs, including FGFR^4,5^, EphB2^6^, and the neurotrophin receptors TrkA, TrkB, and TrkC^7–9^. The blue light-induced clustering of CRY2 lends itself especially well to reproducing the conditions for RTK activation. Previous work has shown that fusion of CRY2 to the C-terminal of either the full length receptor^8^, or a truncated version which lacks the ECD and TM and is trafficked to the membrane via a Lyn myristoylation tag^7,10^, is sufficient to initiate downstream signaling in a light-dependent manner. In our recent work^7^, we evaluated both of these methods of constructing a CRY2 opto-TrkA. In addition, we offered a third design that expresses CRY2 fused to the ICD of TrkA (iTrkA, aa 450-799) with no membrane-targeting sequence: in the dark, CRY2-iTrkA is inactive and soluble in the cytosol. Upon blue light illumination, iTrkA is activated via the clustering of CRY2, and can simultaneously be recruited to the plasma membrane (or other subcellular membranes of interest) via the interaction of CRY2 with CIBN. In our hands, we find that this CRY2-iTrkA construct affords low dark background and robustly reproduces the function of endogenous TrkA upon light stimulation-with a greater efficacy than either our full-length or myristoylated opto-TrkA constructs.

While these results are encouraging and demonstrate the potential of CRY2-based opto-RTKs, there are drawbacks to using CRY2 for optogenetic manipulations. While CRY2 opto-RTKs can reproduce endogenous signaling, they are somewhat artificial in that they form oligomers, rather than dimers, of the RTK of interest. Despite attempts to engineer CRY2, it is still not possible to precisely limit the number of CRY2 molecules present in light-induced clusters, which removes a degree of control. Moreover, a key goal of our work is to limit RTK activation to specific subcellular locations via genetically encoded peptide tags in order to assess the impact of cellular context on signaling outcome. While CIBN can recruit CRY2 clusters to the plasma membrane or other subcellular locations of interest, it is still possible for unbound, cytosolic CRY2 to cluster and activate the opto-RTK simultaneously, confounding the signal output from specific compartments. Finally, another often-cited advantage of using light to activate protein signaling is the temporal precision offered: optogenetic strategies offer fine control of the duration of activation in a way that is not possible when using endogenous ligand. However, while light induces CRY2 clustering and CIBN binding on the order of seconds, complete dissociation requires tens of minutes^3^, which limits the temporal resolution of activation. To address these limitations, we have designed a strategy which uses the improved Light-Induced Dimer (iLID) and a tandem-dimer construct of SspB^3,11^ to achieve location-specific activation of TrkA and TrkB in a light-dependent manner in live cells. We offer this tandem-dimer construct as a generalizable tool for activation of other RTKs with greater control of kinetics^12^, affinity, and localization than other available opto-RTK systems.

## Results

### Design of iLID opto-iTrkA

The binding of a neurotrophin to its receptor, such as NGF to TrkA or BDNF to TrkB, stabilizes the active dimer conformation of the receptor^2^. As CRY2 forms homo-oligomeric clusters in the presence of blue light (**Figure 1a**), we have noted that our previously reported CRY2-iTrkA system^7^ can induce the formation of iTrkA oligomers in the cytosol as well as on the plasma membrane by binding to CIBN (**Figure 1b**) and subsequently activate downstream signaling cascades (**Supplementary Figure 1**). In order to more closely mimic endogenous Trk activation, and to attain better control of the subcellular location of activation, we have designed our iLID opto-iTrk system to form a Trk dimer only upon recruitment to SspB.

**Figure 1:**
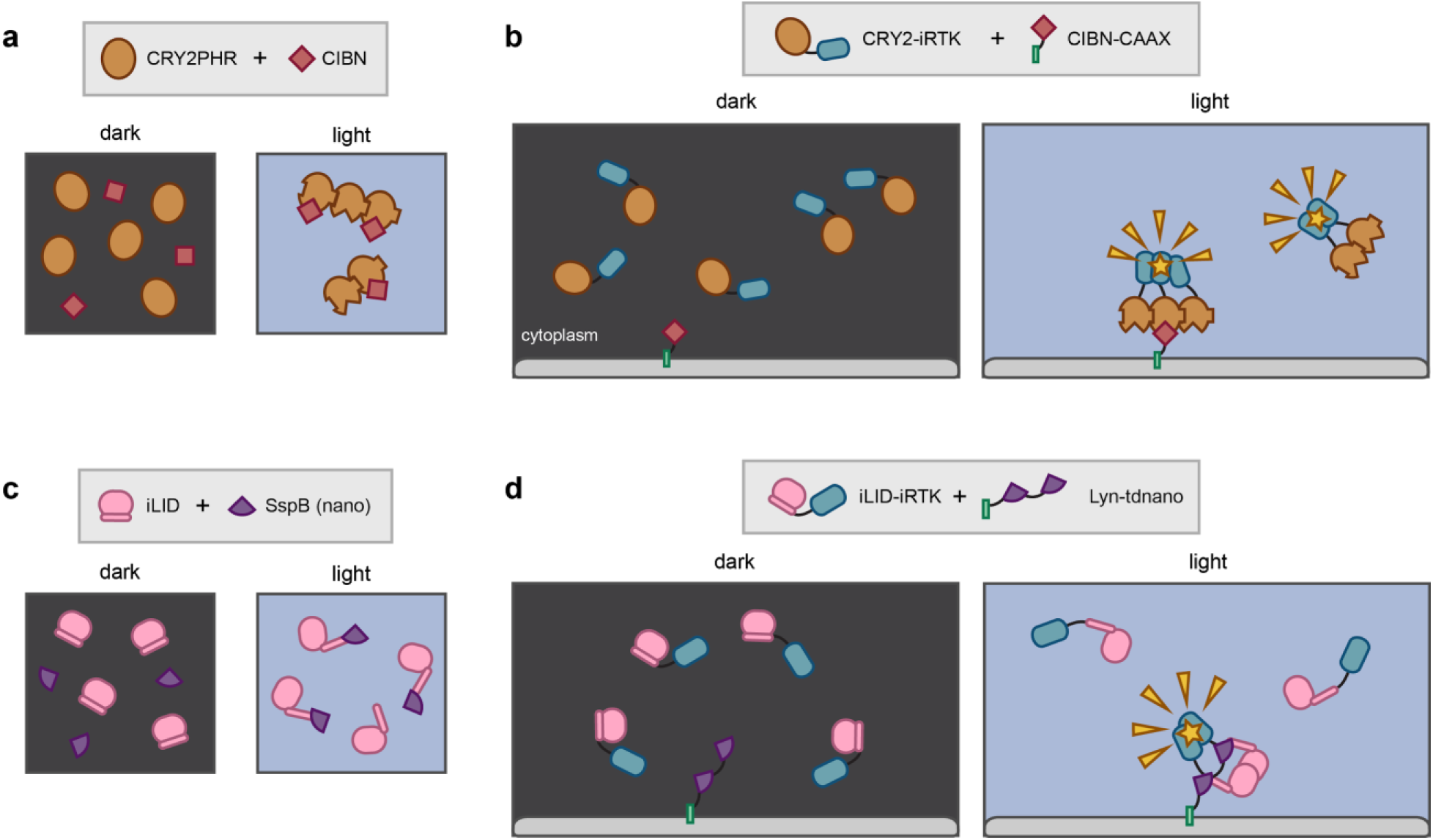
The iLID + tdnano system permits light-activation of RTKs with more precise control than CRY2. **a**) In response to blue light stimulation, CRY2PHR is known to form homo-oligomeric clusters in addition to binding its partner CIBN. **b**) The design of a system using a cytosolic CRY2-RTK and a membrane bound CIBN. While location-specificity can be achieved by a genetic tag on CIBN, higher-order RTK oligomers can form and become active in the cytosol or other subcellular locations. **c**) iLID and its partner SspB, or nano, form a 1 to 1 heterodimer, with no higher-order clustering. **d**) By genetically targeting a tandem dimer of SspB, dubbed tdnano, to a specific membrane, we achieve formation of an active RTK dimer exclusively at that region of interest.

The design of iLID incorporates the small peptide SsrA into the J-alpha helix of the *Avena sativa* LOV2 domain^11^. In the dark, this helix is caged and SsrA is unavailable for interaction with its partner SspB (the wild type SspB has been dubbed nano by Kuhlman, referring to its nanomolar affinity for iLID). Upon light illumination, the J-alpha helix unfolds, permitting SsrA-SspB binding. The light-induced conformational change of iLID permits the formation of a one-to-one heterodimer (**Figure 1c**). In order to induce the formation of an iTrk dimer upon blue light stimulation, we have designed a tandem-dimer construct with two copies of nano bridged by a long, flexible (GGSGGSGGSGGSGGGS) linker, dubbed tdnano (**Figure 1d**). We target this tandem dimer to the plasma membrane via a genetically-encoded Lyn myristoylation tag. Unlike in the CRY2 opto-iTrkA system, iLID-RTK alone does not oligomerize upon blue light stimulation and should not initiate downstream signaling: the iLID opto-iTrk design requires the presence of tdnano for dimerization, and thus for activation, increasing our confidence that any signaling triggered by light is coming from the subcellular location of interest. It is worth noting that in our hands, using N-terminal targeting sequences for tdnano proved robust, while tdnano bearing very short C-terminal targeting sequences such as CAAX appeared to mis-localize (data not shown).

### iLID opto-iTrkA allows real-time kinetic studies of downstream signaling in live cells

In order to image both the fluorescent activity reporter and the opto-iTrkA construct simultaneously, we sacrifice labeling tdnano with a fluorescent protein, and instead affix a small, triple-HA epitope tag. We have confirmed by confocal microscopy on immunostained samples that Lyn-tdnano expressed in PC12 is located primarily at the plasma membrane, with some additional staining on intracellular vesicles (**Supplementary Figure 2**). Further, to ensure that our opto-iTrks are binding to Lyn-tdnano, we performed a translocation test in PC12 cells. Though subtle, we observe a rapid decrease in intensity of mCherry expression in the cytosol of co-transfected cells, suggesting successful recruitment of the iLID opto-iTrk to the membrane via blue light-dependent interaction with Lyn-tdnano (**Supplementary Figure 3**).

**Figure 2:**
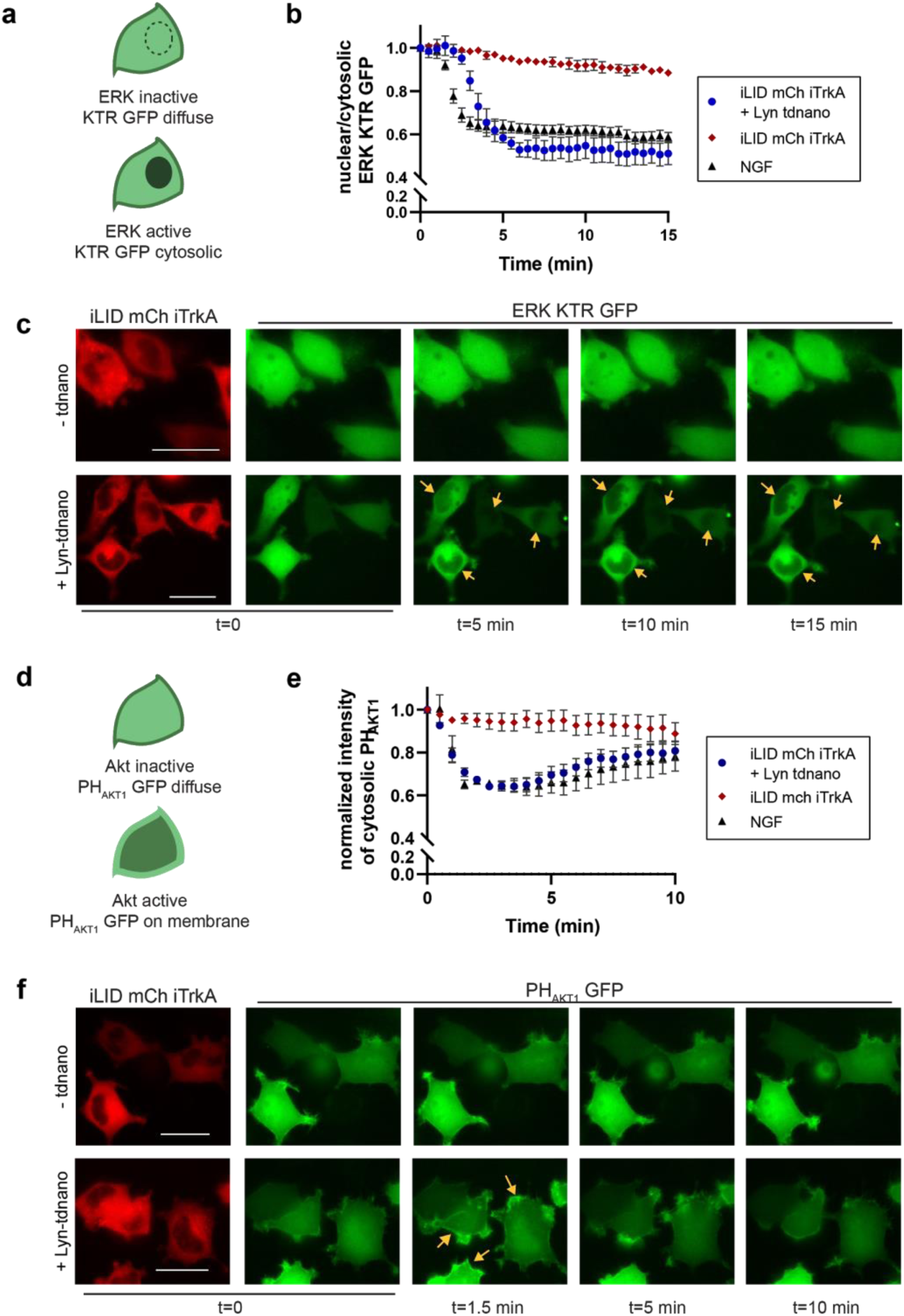
iLID-iTrkA in the presence of Lyn-tdnano activates pathways downstream of NGF/TrkA. **a**) When ERK is inactive, the ERK kinase translocation reporter (KTR) is diffuse through the whole cell. Active ERK phosphorylates the KTR, which leads to it being shuttled out of the nucleus. **b**) ERK activity quantified as nuclear/cytosolic ratio of ERK KTR GFP. iLID-iTrkA alone is insufficient to activate ERK. Co-expression of iLID-iTrkA with Lyn-tdnano leads to ERK activation at the plasma membrane upon blue light stimulation (n=4 cells/condition). **c**) Representative images of ERK KTR translocation experiments. **d**) When Akt is inactive, the pleckstrin homology domain of Akt1 (PH_AKT1_) shows diffuse expression in the cell. Upon Akt activation, PH_AKT1_ translocates to the plasma membrane. **e**) Quantification of Akt activity as decrease in cytosolic intensity of PH_AKT1_. With blue light stimulation, expression of iLID-iTrkA alone does not show Akt activation, but coexpression of iLID-iTrkA with Lyn-tdnano shows similar activation of Akt as treatment with NGF (n=6 cells/condition). **f**) Representative images of PH_AKT1_ translocation experiments. Scale bars = 20 μm. NGF treatment performed at concentration of 50 ng/mL.

**Figure 3:**
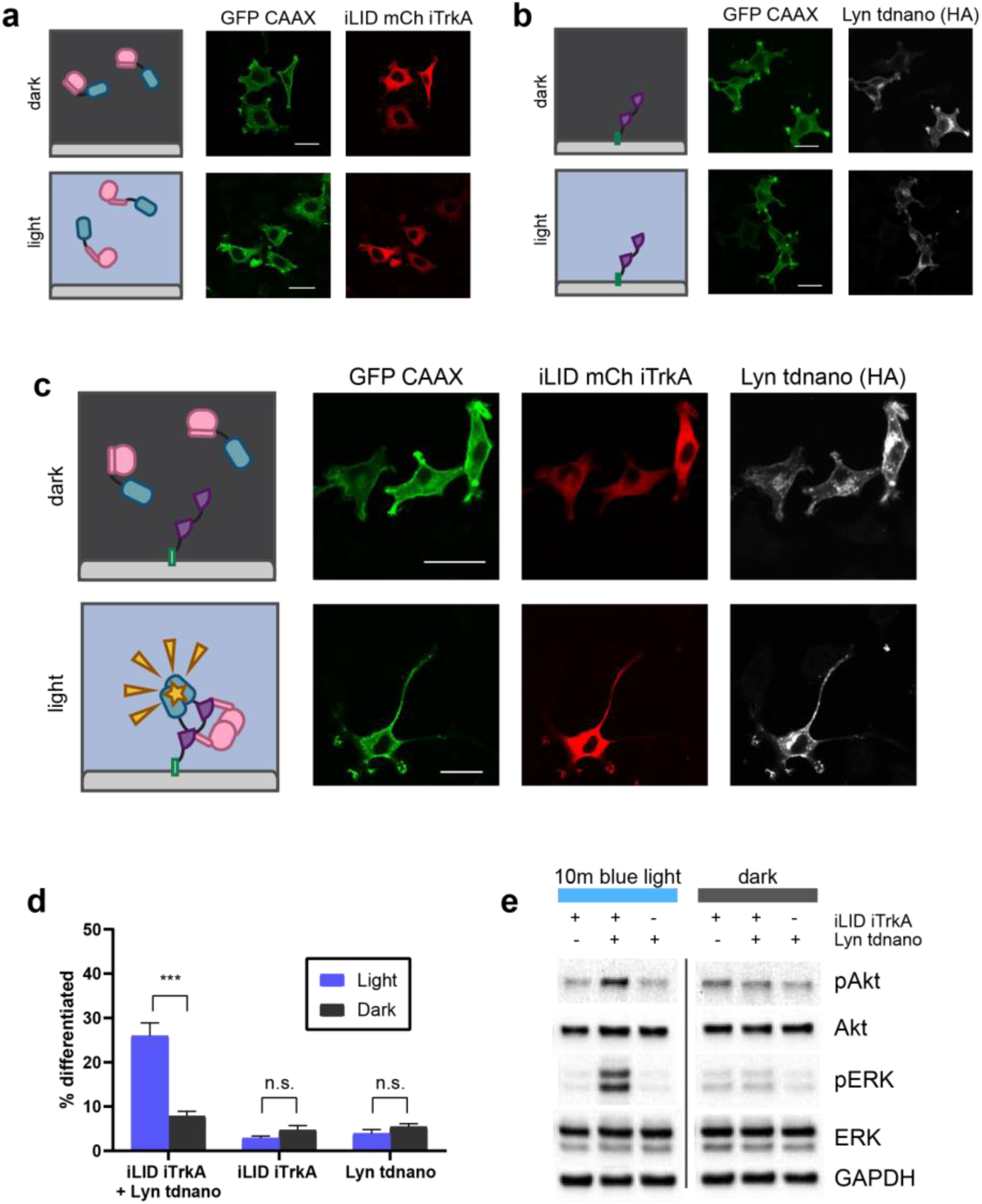
iLID-iTrkA permits both multi-day and population-level studies. iLID-iTrkA (**a**) or Lyn-tdnano (**b**) alone are insufficient to induce neurite growth in PC12 cells, either with or without blue light illumination.. **c**) Coexpression of iLID-iTrkA with Lyn-tdnano shows a robust increase in PC12 cell differentiation when cells are illuminated with blue light (5 min on/5 min off, 400 μW/cm^2^ for 40h). **d**) Quantification of neurite growth experiments: cells were counted as differentiated if they grew neurites >20 μm long. (n>90 cells per trial per condition with 3 independent trials) **e**) Immunoblot from transfected PC12 cells, treated with 10 min blue light or left in the dark. Phosphorylation of ERK and Akt is only observed when both iLID-iTrkA and Lyn-tdnano are expressed. Scale bars = 20 μm.

We next assayed the activation of signaling pathways downstream of TrkA using live cell reporter assays. Upon activation by neurotrophin binding, TrkA in turn activates several signaling cascades, including the MAPK/ERK and PI3K/Akt pathways^13^. To visualize the activity of these pathways in live cells in real-time, we made use of an ERK Kinase Translocation Reporter^14^ (KTR) fused to GFP and the pleckstrin homology domain of Akt1^15^ (PH_AKT1_) fused to GFP, respectively.

The ERK KTR was designed to bind and become phosphorylated by active ERK. In the absence of ERK signaling, the reporter is visible in the whole cell. Upon phosphorylation by active ERK, the reporter moieties in the nucleus translocate to the cytosol (**Figure 2a**). We observe that when iLID opto-iTrkA is expressed without the nano tandem dimer, there is no translocation of the ERK KTR. However, when co-expressed with Lyn-tdnano, we observe robust translocation of KTR, thereby signaling activation of ERK, within 5 minutes of initial blue light stimulation (**Figure 2b, c**).

The activation of AKT signaling was assayed using the PH_AKT1_ reporter. When Akt is phosphorylated, PH_AKT1_ translocates from the cytosol to the plasma membrane (**Figure 2d**). When iLID-iTrkA is expressed alone in PC12 cells without the tdnano construct, translocation of PH_AKT1_ is not observed upon illumination with blue light. When the opto-iTrks are co-expressed with the membrane-bound Lyn-tdnano, however, we observe transient translocation of PH_AKT1_ approximately 90 seconds after beginning light stimulation, with kinetics similar to those observed upon NGF treatment (**Figure 2e, f**).

### iLID opto-iTrkA is suitable for population-level and multi-day studies

In addition to live-cell experiments, we sought to demonstrate the compatibility of iLID opto-iTrks with multi-day experiments. To this end, we performed a neurite growth assay using iLID opto-iTrkA. When PC12 cells are grown in the presence of NGF, they differentiate and assume a “neuron-like” morphology, featuring the growth of long neurites. This morphological change occurs over just 2-3 days in the NeuroScreen1 subclone used in our lab. We transfected cells with either both components of our opto-iTrkA system (iLID-iTrkA + Lyn-tdnano) or with a single component (iLID-iTrkA alone or Lyn-tdnano alone) and allowed cells to recover for 24 hours. We then changed the cells to minimal-serum media and either illuminated for 40 hours with intermittent light (5 min on, 5 min off), or left cells in the dark for 40 hours. Cells were fixed, stained to visualize Lyn-tdnano expression, and imaged. Cells exhibiting fluorescence corresponding to transfected constructs were counted, and were considered differentiated if they bore neurites greater than 20 um in length. The percent of differentiated cells in each condition, as well as representative images from each condition, are shown in **Figure 3**. Cells grown in the dark showed <10% differentiation, and the percent of differentiated cells did not increase significantly with light when either the iLID-iTrkA or Lyn-tdnano constructs were expressed alone. However, the percent of differentiated cells was significantly different between light and dark conditions when both components of the opto-iTrkA system were expressed (26.0 ± 5.0% with light vs 7.8 ± 1.9% in the dark).

We next sought to demonstrate the applicability of iLID opto-iTrkA at the population level. To this end, we used a home-built LED array to illuminate cells in culture. PC12 cells were transfected with either iLID-iTrkA alone, Lyn-tdnano alone, or both iLID-iTrkA and Lyn-tdnano. After recovery, cells were serum-starved to reduce basal signaling, and then illuminated with low-power (400 uW) blue light for 10 minutes or left in the dark. Cells left in the dark show consistently low levels of phosphorylation of ERK and Akt, as do illuminated populations that express only one of the two components of our opto-iTrk tools. However, cells that express both iLID-iTrk and Lyn-tdnano show a dramatic increase in phosphorylation of ERK and Akt downstream of opto-iTrk signaling in the presence of light, with no significant activity in the dark (**Figure 3e**). These results together with the neurite growth data suggest that the iLID opto-iTrk tools are robust enough to allow studies of signaling dynamics on the population-scale and over long time in addition to short-term, single-cell studies.

### iLID opto-iTrkB demonstrates that Lyn-tdnano can be generalized to construct other opto-RTKs

In addition to our specific interest in studying the signaling of TrkA, we wanted to demonstrate that the iLID + tdnano system is generalizable to other receptor tyrosine kinases of interest. As a proof of concept, we constructed an iLID opto-iTrkB utilizing the intracellular signaling domain of human TrkB (aa 455-822). We characterized this construct in live cells studies, again using the ERK KTR and PH_AKT1_ reporters to assess the activity of downstream pathways. As with opto-iTrkA, we did not observe any activation of downstream signaling pathways with iLID-iTrkB was expressed alone. However, when co-expressed with Lyn-tdnano, we saw robust activation of both ERK and Akt (**Figure 4**). It is especially interesting to note that iLID-iTrkB activation appeared to induce growth cone formation, visible via the membrane-localized PH_AKT1_-GFP. The ability to induce new growth cone and neurite formation has been observed by others in the context of CRY2-based opto-TrkB systems^16^.

**Figure 4:**
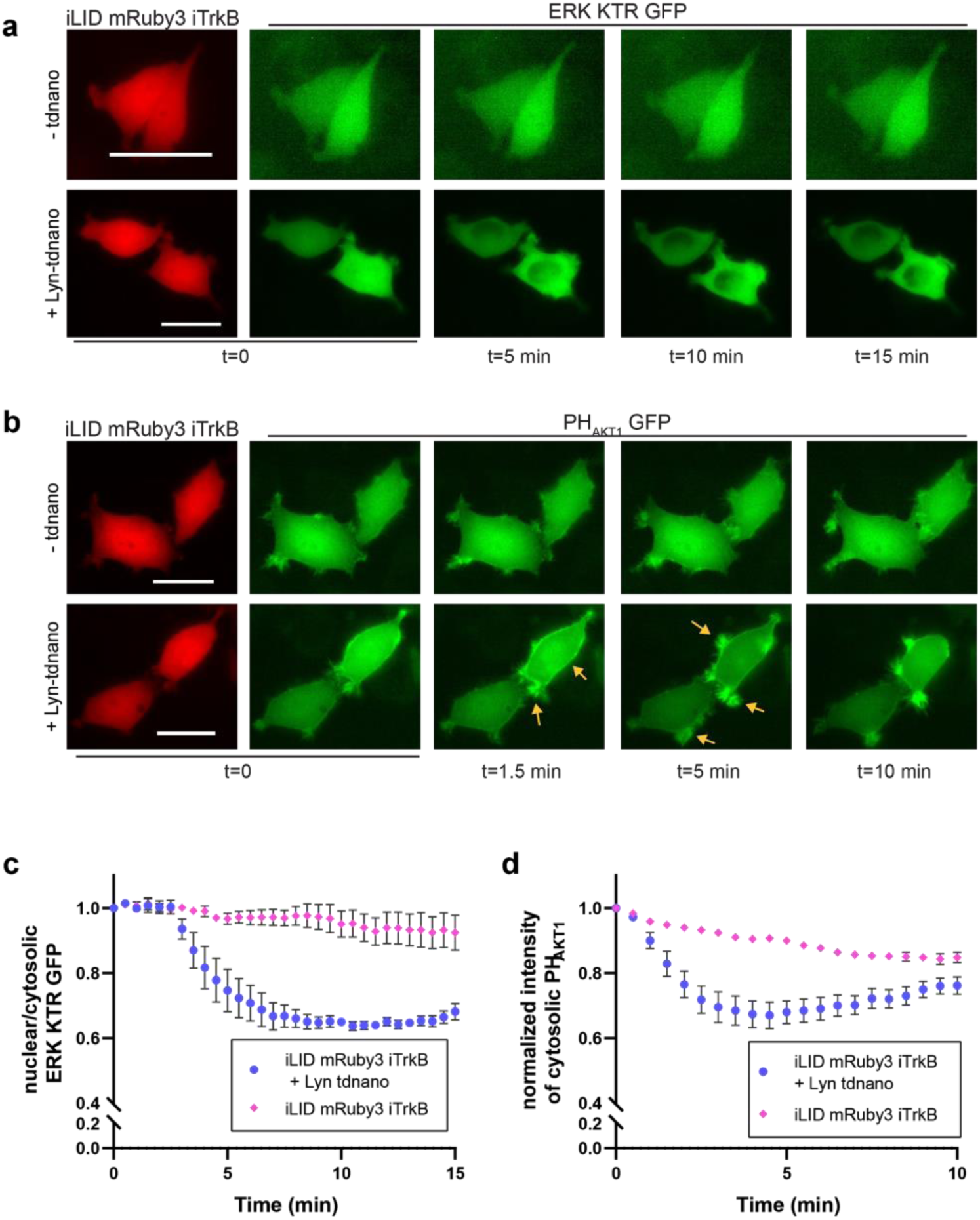
We demonstrate that the iLID-iRTK + Lyn-tdnano strategy is generalizable to other receptors of interest using iLID opto-iTrkB. **a**) Representative timecourse of ERK KTR nuclear export assay with iLID-iTrkB alone or with iLID-iTrkB + Lyn-tdnano. **b**) Representative timecourse of PH_AKT1_ membrane translocation assay with iLID-iTrkB alone or with iLID-iTrkB + Lyn-tdnano. When both components are expressed, PH_AKT1_ is enriched lamellipodia. **c**) Quantification of ERK activity as nuclear/cytosolic ratio of ERK KTR GFP. Expression of iLID-iTrkB alone does not show ERK activation, but coexpression with Lyn-tdnano exhibits sustained ERK activation (n=5-6 cells/condition). **d**) Quantification of Akt activity as decrease in cytosolic intensity of PH_AKT1_. Expression of iLID-iTrkB alone does not show Akt activation, but coexpression with Lyn-tdnano displays robust activation of Akt (n=5-6 cells/condition). Scale bars = 20 μm.

## Discussion

In this work we aim to demonstrate the unique utility of an iLID-based optogenetic method for the activation of receptor tyrosine kinases. Kuhlman’s iLID-nano dimerizer has proven powerful in offering optogenetic control of cellular processes via reconstitution of split proteins^17,18^ or by using the formation of the heterodimer to recruit a protein of interest to a particular subcellular location^19^. In this work, by constructing a tandem dimer nano construct, we utilize the tool to simultaneously dimerize and alter the localization of a molecule of interest, a method that is especially useful for those seeking fine spatial and temporal control of RTK signaling. We have demonstrated its ability to activate two widely studied receptor tyrosine kinases, TrkA and TrkB, and we believe the tool has the potential to be more broadly generalized to other classes of RTKs beyond these neurotrophin receptors.

Light-gated receptor tyrosine kinases have been designed using several different strategies, and existing opto-RTKs are very powerful in many applications. Other groups have successfully built opto-TrkA or opto-TrkB systems that are permanently tethered to the plasma membrane by the Lyn myristoylation motif, using CRY2^16^, which clusters under blue light, or AuLOV^10^, which dimerizes in response to blue light. However, by permanently tethering the signaling domain of TrkA to the plasma membrane, one increases the local concentration of the receptor. Especially when using transient transfection to deliver constructs to cells, this raises the issue that overexpression coupled with higher local concentrations can lead to high basal activity, even in the absence of light. In our design, the signaling domain is dissociated from the membrane until activation is desired. Another recent work similarly decoupled the activation from the localization, by making the dimerization of TrkA dependent on far-red light and the recruitment to the membrane blue-light dependent^20^. While this system can offer that extra degree of control, it requires two sets of photoreceptors, adding to the complexity. Finally, the iLID-SspB system is especially appealing as it has been continually updated, both in tuning its off kinetics, with the “slow” sLID^12^, and in tuning the binding affinity of the dimer pair with the stronger binding micro and milli mutants of SspB^11,12^. We believe that using iLID with a tandem-dimer of SspB allows for finer control of organelle-specific signaling without increasing the complexity of the system, and the Lyn motif can be easily replaced with other N-terminal signal peptides or domains for targeting to other regions of interest.

## Methods

### Plasmid construction

pLL7.0: Venus-iLID-CAAX (Addgene plasmid #60411) and pLL7.0: tgRFPt-SSPB WT (Addgene plasmid #60415) were gifts from Brian Kuhlman^11^. To generate iLID-mCherry-iTrkA, the iLID-encoding sequence was PCR-amplified from Venus-iLID-CAAX and inserted into CRY2PHR-mCh-iTrkA^7^ at XhoI and SmaI using In-Fusion (Clontech). iLID-mRuby3-iTrkB was generated by PCR amplification of iLiD, mRuby3 and iTrkB (aa 455-822) from separate vectors and inserted into a pmCherry-N1 vector at EcoRI and NotI using In-Fusion. To generate Lyn-nano-nano-3xHA, separate gblocks were synthesized (IDT) encoding Lyn-SspB WT, SspB WT, and a 3xHA tag, then assembled and inserted into pEGFP-N1 at NheI and NotI using In-Fusion (Clontech). These plasmids have been deposited with Addgene.

### Cell culture and transfection

PC12 cells (NeuroScreen-1 subclone, Cellomics, discontinued) were maintained in complete medium (F12K supplemented with 15% horse serum and 2.5% FBS, all from Thermo Fisher) and grown in a standard incubator at 37°C with 5% CO2. Cells tested negative for mycoplasma contamination by PCR with primers 5’-GTGGGGAGCAAAYAGGATTAGA-3’ and 5’-GGCATGATGATTTGACGTCRT-3’^21^.

For live cell kinase activity assays, PC12 cells were transfected using Turbofect (Thermo Fisher Scientific) according to the manufacturer’s protocol. The transfected cells were allowed to recover and express desired constructs overnight in complete culture medium. Cells were serum-starved (F12K with 1.5% horse serum and 0.25% FBS) for 4-6 hours prior to imaging to reduce basal kinase activity.

For neurite growth and immunoblotting assays, PC12 cells were transfected by electroporation using the Amaxa Nucleofector II (Lonza). Cells were added to a suspension of DNA in electroporation buffer (7 mM ATP, 11.7 mM MgCl_2_, 86 mM KH_2_PO_4_, 13.7 mM NaHCO_3_, 1.9 mM glucose), transferred to a 2mm electroporation cuvette (Fisher Scientific) and subjected to the manufacturer provided protocol for PC12 cells, then plated on poly-L-lysine coated coverslips or culture dishes. Cells were allowed to recover for 40 h in complete medium, then serum-starved for 8 h prior to experiment.

### Live cell imaging: kinase activation assays

Live cell imaging was performed on an epifluorescence microscope (Leica DMI6000B) equipped with an on-stage CO_2_ incubation chamber (Tokai Hit GM-8000) and a motorized stage (Prior). An adaptive focus control was used to actively keep the image in focus during the period of imaging. A light-emitting diode light engine (Lumencor) was used as the light source for fluorescence imaging. Pulsed blue light (200 ms pulse duration at 9.7 W/cm2) was delivered every 10 s both to image GFP-tagged constructs and to initiate iLID/SspB interactions. Pulsed green light (200 ms pulse duration) was used to image mCherry. The microscope was equipped with a commercial GFP filter cube (Leica; excitation filter 472/30, dichroic mirror 495, emission filter 520/35) and a commercial Texas Red filter cube (Leica; excitation filter 560/40, dichroic mirror 595, emission filter 645/75). Images acquired with an oil-immersion 100× objective and imaged with a sensitive CMOS camera (PCO.EDGE 5.5) (PCO). ERK KTR translocation assays were quantified by measuring GFP intensity of the whole nucleus and whole cell in ImageJ, then plotted as nuclear/cytosolic intensity over time, normalized to the nuclear/cytosolic intensity at t_0_. PH_AKT1_ translocation assays were quantified by measuring the average GFP intensity of the cytosol, and plotted over time, normalized to the cytosolic intensity at t_0_.

### Programmable LED light box

LED boxes were constructed as previously reported for long-term light illumination.^22^ Briefly, a 12 well plate sized blue LED array was constructed by assembling 12 blue LEDs (LED465E, ThorLabs) on a breadboard, housed in an aluminum box, and a light diffuser film was positioned above the LED array to ensure uniform light intensity in defined area. The light intensity was measured by a power meter (Newport, 1931-C). The LED array on/off timing was controlled by an Arduino UNO.

### Immunoblotting

After desired treatment, cultured cells were moved to ice, rinsed with ice cold PBS, and lysed in RIPA buffer (25 mM Tris HCl, 150mM NaCl, 1% Triton X-100, 1% Sodium deoxycholate, 0.1% SDS) supplemented with protease and phosphatase inhibitor cocktails (Roche 04906837001 and 04693132001). Clarified lysates were mixed with Laemmli sample buffer (Bio-Rad 1610747) and β-mercaptoethanol and boiled for 10 min. Lysed samples were subjected to electrophoresis using Bio-Rad’s Mini-PROTEAN system (1658026FC). After separation, protein was transferred to a nitrocellulose membrane (Bio-Rad), followed by standard blotting procedure. Primary antibodies were obtained from Cell Signaling Technology: anti-pAKT (T308) (CST 9275), anti-AKT (CST 9272), anti-pERK1/2 (T202+Y204) (CST 9101), anti-ERK1/2 (CST 9102), and anti-GAPDH (CST 2118). HRP-conjugated secondary antibody (CST 7074) was used for protein band detection. Protein bands were visualized by chemiluminescence (Bio Rad 1705060) using a ChemiDoc imaging system.

### Neurite growth assay

For neurite growth experiments, transfected PC12 cells were kept in dark and serum-starved for 8 h prior to blue light stimulation to minimize interference from growth factors present in serum. Cells were then illuminated with blue light under a 5 min on/5 min off protocol at 400 μW/cm^2^ for 40 h using a custom-built LED array housed inside a CO2 incubator. This illumination condition has been repeatedly tested and shows no obvious toxicity as demonstrated in our previous studies.^22,23^ Additional sets of transfected cells were kept in dark as controls. After 40 h, cultures were fixed using 4% paraformaldehyde. The HA-tagged tdnano construct was stained using a rat α-HA primary antibody (Roche) and an Alexa Fluor 647-conjugated secondary antibody (Thermo Fisher). Samples were imaged using a Nikon A1R confocal microscope with an oil-immersion 63× objective using 488 nm, 561 nm, and 647 nm lasers.

## Acknowledgements

We thank Dr. Tobias Meyer (Stanford University) for providing PH_Akt1_-GFP and the PC12 NeuroScreen1 cell line, and Dr. Markus Covert (Stanford University) for providing ERK KTR. This work was supported by the US NIH (DP2-NS082125) and a National Science Foundation Graduate Research Fellowship under Grant No. DGE-1656518 (JMH).

---

**Supplementary Figure 1:**
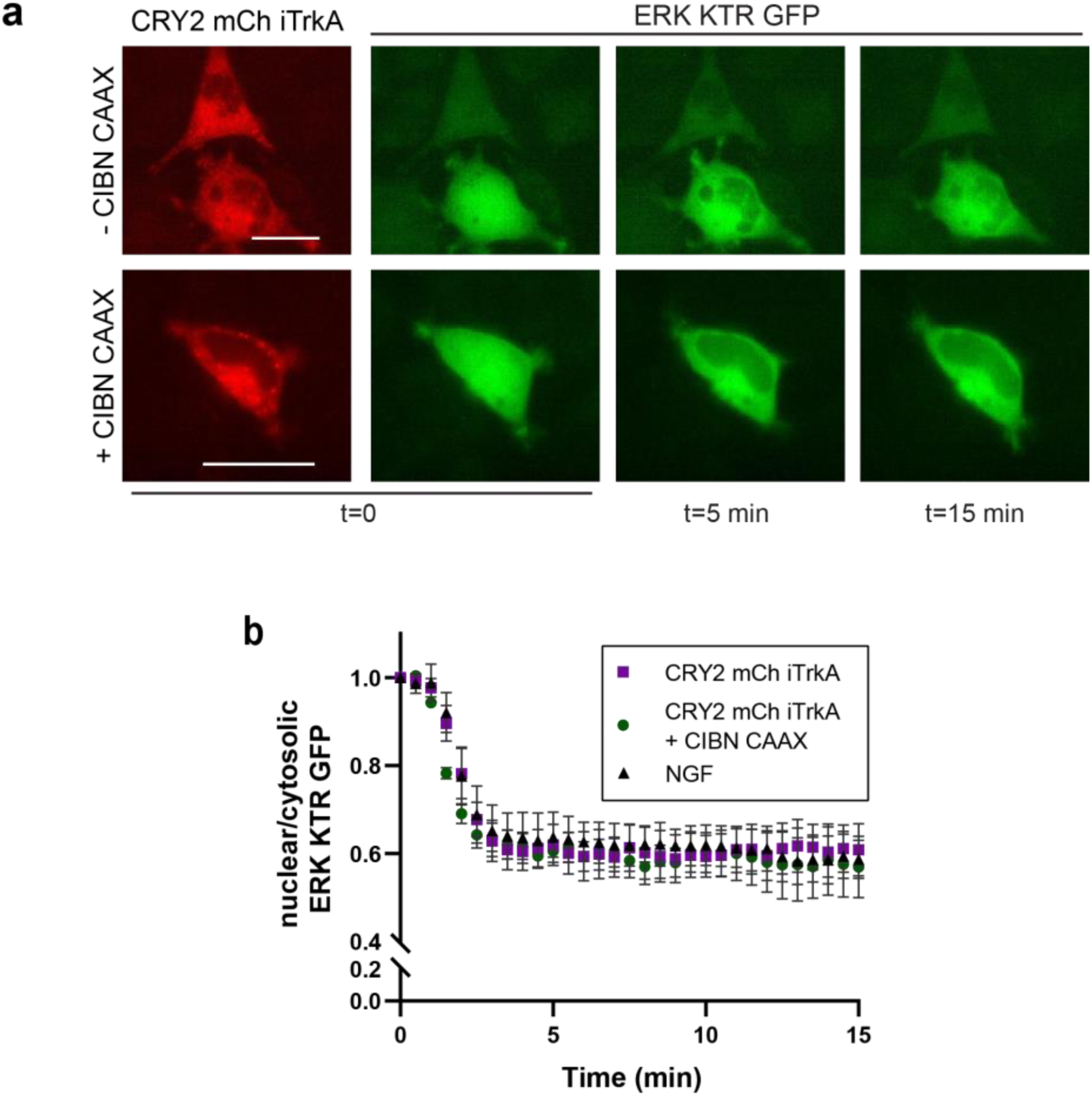
CRY2 opto-iTrkA is active both with and without the co-expression of CIBN-CAAX. **a**) Representative timecourse of ERK KTR GFP translocation with CRY2-iTrkA alone (top) or CRY2-iTrkA + CIBN CAAX (bottom). **b**) Quantification of ERK activity as normalized nuclear/cytosolic ERK KTR GFP (n=4-6 cells per condition). Scale bars = 20 μm.

**Supplementary Figure 2:**
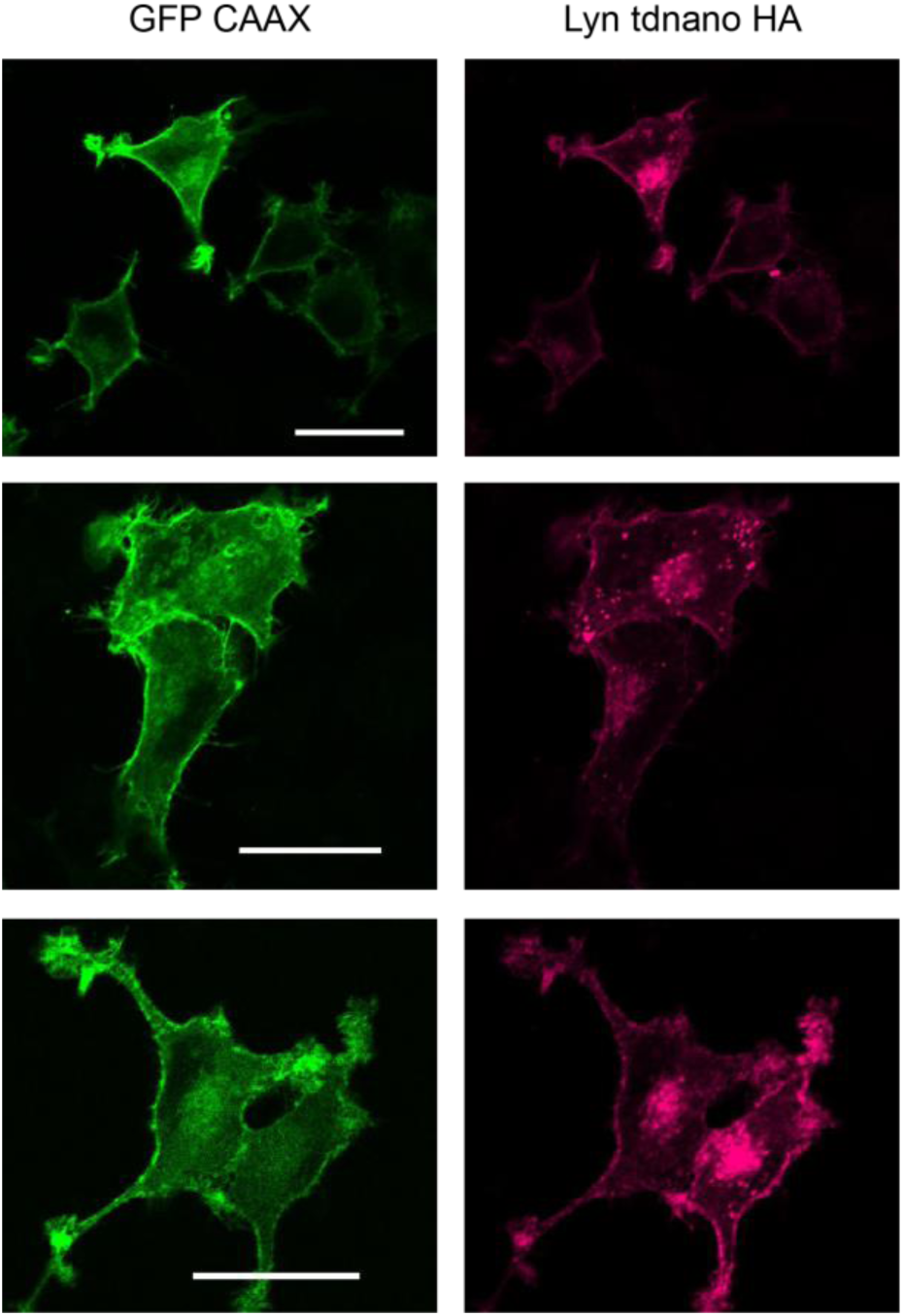
Lyn-tdnano expressed in PC12 cells localizes to the plasma membrane, with some presence on internal membrane structures. Cell membrane is labeled with GFP CAAX (left), and Lyn-tdnano is detected by immunostaining for HA. Images obtained with confocal microscopy. Scale bars = 20 μm.

**Supplementary Figure 3:**
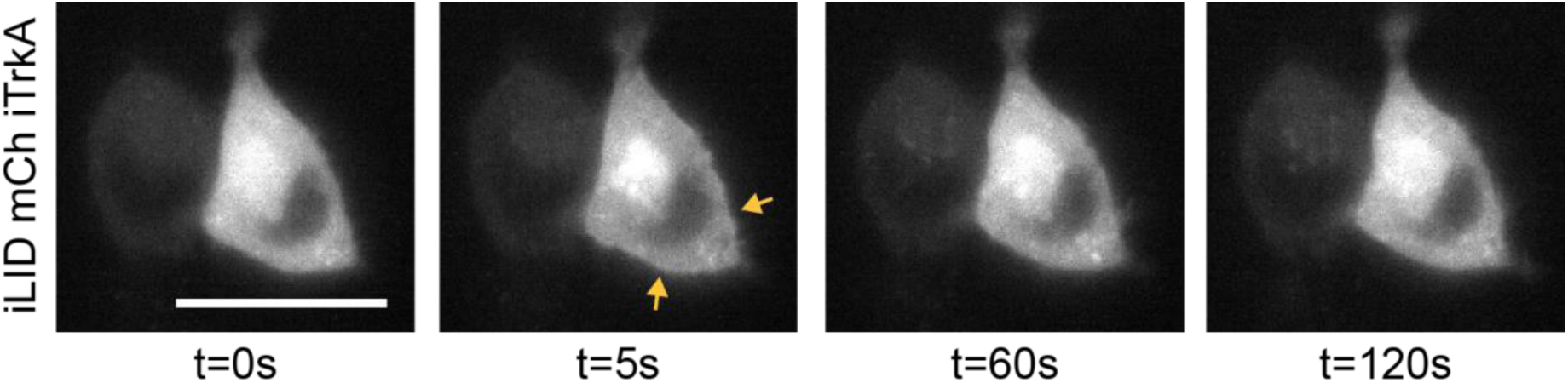
iLID-iTrkA in the presence of Lyn-tdnano translocates rapidly. Prior to blue light illumination, iLID-iTrkA is diffuse in the cytosol. Following a single 2s pulse of blue light, the construct is recruited to the plasma membrane as well as internal membrane structures. This recruitment is almost entirely reversed after only two minutes in the dark. Images acquired with epifluorescent microscopy. Scale bar = 20 μm.

## References

1. Ullrich, A. & Schlessinger, J. Signal transduction by receptors with tyrosine kinase activity. Cell 61, 203–212 (1990).

2. Lemmon, M. A. & Schlessinger, J. Cell signaling by receptor tyrosine kinases. Cell 141, 1117–1134 (2010).

3. Kennedy, M. J. et al. Rapid blue-light-mediated induction of protein interactions in living cells. Nat. Methods 7, 973–975 (2010).

4. Kim, N. et al. Spatiotemporal control of fibroblast growth factor receptor signals by blue light. Chem. Biol. 21, 903–912 (2014).

5. Bugaj, L. J. et al. Regulation of endogenous transmembrane receptors through optogenetic Cry2 clustering. Nat. Commun. 6, 6898 (2015).

6. Locke, C., Machida, K., Tucker, C. L., Wu, Y. & Yu, J. Optogenetic activation of EphB2 receptor in dendrites induced actin polymerization by activating Arg kinase. Biology Open 6, 1820–1830 (2017).

7. Duan, L. et al. Optical Activation of TrkA Signaling. ACS Synthetic Biology 7, 1685–1693 (2018).

8. Chang, K.-Y. et al. Light-inducible receptor tyrosine kinases that regulate neurotrophin signalling. Nature Communications 5, (2014).

9. Han, K. A. et al. Neurotrophin-3 Regulates Synapse Development by Modulating TrkC-PTP Synaptic Adhesion and Intracellular Signaling Pathways. Journal of Neuroscience 36, 4816–4831 (2016).

10. Khamo, J. S., Krishnamurthy, V. V., Chen, Q., Diao, J. & Zhang, K. Optogenetic Delineation of Receptor Tyrosine Kinase Subcircuits in PC12 Cell Differentiation. Cell Chem Biol 26, 400–410.e3 (2019).

11. Guntas, G. et al. Engineering an improved light-induced dimer (iLID) for controlling the localization and activity of signaling proteins. Proc. Natl. Acad. Sci. U. S. A. 112, 112–117 (2015).

12. Zimmerman, S. P. et al. Tuning the Binding Affinities and Reversion Kinetics of a Light Inducible Dimer Allows Control of Transmembrane Protein Localization. Biochemistry 55, 5264–5271 (2016).

13. Sofroniew, M. V., Howe, C. L. & Mobley, W. C. Nerve growth factor signaling, neuroprotection, and neural repair. Annu. Rev. Neurosci. 24, 1217–1281 (2001).

14. Regot, S., Hughey, J. J., Bajar, B. T., Carrasco, S. & Covert, M. W. High-sensitivity measurements of multiple kinase activities in live single cells. Cell 157, 1724–1734 (2014).

15. Raucher, D. et al. Phosphatidylinositol 4,5-Bisphosphate Functions as a Second Messenger that Regulates Cytoskeleton–Plasma Membrane Adhesion. Cell 100, 221–228 (2000).

16. Woo, D. et al. Locally Activating TrkB Receptor Generates Actin Waves and Specifies Axonal Fate. Cell Chem Biol (2019). doi :10.1016/j.chembiol.2019.10.006

17. Liu, Q. et al. A Photoactivatable Botulinum Neurotoxin for Inducible Control of Neurotransmission. Neuron 101, 863–875.e6 (2019).

18. Pu, J., Zinkus-Boltz, J. & Dickinson, B. C. Evolution of a split RNA polymerase as a versatile biosensor platform. Nat. Chem. Biol. 13, 432–438 (2017).

19. O’Neill, P. R., Kalyanaraman, V. & Gautam, N. Subcellular optogenetic activation of Cdc42 controls local and distal signaling to drive immune cell migration. Mol. Biol. Cell 27, 1442–1450 (2016).

20. Leopold, A. V., Chernov, K. G., Shemetov, A. A. & Verkhusha, V. V. Neurotrophin receptor tyrosine kinases regulated with near-infrared light. Nat. Commun. 10, 1129 (2019).

21. Molla Kazemiha, V. et al. PCR-based detection and eradication of mycoplasmal infections from various mammalian cell lines: a local experience. Cytotechnology 61, 117–124 (2009).

22. Zhang, K. et al. Light-mediated kinetic control reveals the temporal effect of the Raf/MEK/ERK pathway in PC12 cell neurite outgrowth. PLoS One 9, e92917 (2014).

23. Che, D. L., Duan, L., Zhang, K. & Cui, B. The Dual Characteristics of Light-Induced Cryptochrome 2, Homo-oligomerization and Heterodimerization, for Optogenetic Manipulation in Mammalian Cells. ACS Synth. Biol. 4, 1124–1135 (2015).

